# 2-Photon imaging of fluorescent proteins in living swine

**DOI:** 10.1101/2023.02.14.528533

**Authors:** Beth A. Costine-Bartell, Luis Martinez-Ramirez, Kieran Normoyle, Tawny Stinson, Kevin J. Staley, Kyle P. Lillis

## Abstract

A common point of failure in translation of preclinical neurological research to successful clinical trials comes in the giant leap from rodent models to humans. Non-human primates are phylogenetically close to humans, but cost and ethical considerations prohibit their widespread usage in preclinical trials. Swine have large, gyrencencephalic brains, which are biofidelic to human brains. Their classification as livestock makes them a readily accessible model organism. However, their size has precluded experiments involving intravital imaging with cellular resolution. Here, we present a suite of techniques and tools for *in vivo* imaging of porcine brains with subcellular resolution. Specifically, we describe surgical techniques for implanting a synthetic, flexible, transparent dural window for chronic optical access to the neocortex. We detail optimized parameters and methods for injecting adeno-associated virus vectors through the cranial imaging window to express fluorescent proteins. We introduce a large-animal 2-photon microscope that was constructed with off-the shelf components, has a gantry design capable of accommodating animals > 80 kg, and is equipped with a high-speed digitizer for digital fluorescence lifetime imaging. Finally, we delineate strategies developed to mitigate the substantial motion artifact that complicates high resolution imaging in large animals, including heartbeat-triggered high-speed image stack acquisition. The effectiveness of this approach is demonstrated in sample images acquired from pigs transduced with the chloride-sensitive fluorescent protein SuperClomeleon.

## INTRODUCTION

For studies of neurological disease, useful experimental platforms are available across a spectrum of increasingly complex models ranging from cell culture to invertebrates to rodents to non-human primate (NHP) and humans. However, the clinical translation of findings from extremely reduced preparations directly to human clinical trials commonly fails^1, 2^. In part, this may be due to a lack of proper methodological reporting^3^. Funding bodies have been working to improve translation for over a decade by requiring enhanced focus on reproducibility and rigor^4^. Another strategy for improving translational success for neurological disease therapies involves testing preclinical findings from lower species in large animal models. While testing potential therapies in NHPs is largely avoided due to ethical considerations and cost, swine are large animals that have large, gyrencephalic brains, but are relatively inexpensive compared to NHPs. Swine are raised for food production garnering less ethical concern for their role in biomedical research.

Swine are potentially under-utilized in neurological research, in part because experimental methodologies have not been widely developed. For example, although in vivo viral transduction and 2-photon (2P) imaging has been performed in rodents and NHPs^5-7^, to our knowledge, this technique has yet not been adapted to swine. Not all techniques developed in NHP are immediately adaptable to swine. For example, NHPs tolerate head mounted equipment such as exteriorized imaging ports well; however, head mounted posts are liable to be damaged and/or infected because swine are prone to rub their head against objects. Other challenges to overcome in swine research include the large body size of swine, which presents a greater challenge for movement artifact during imaging. In addition, the skull of swine has much greater accretion and remodeling over time than rodents or NHP^8^. Thus, the dynamic properties of the skull of swine and the position of the brain in relation to the skull must be planned for in long-term studies involving viral transduction and 2P imaging.

Here we describe a “team science” approach, combining neurosurgical, veterinary, and bioengineering expertise to accomplish vivo 2P imaging in swine. Imaging was accomplished in animals up to 84 kg, perhaps the largest subject of any experimental species to date. We describe the use of cortical transfection to express SuperClomeleon and jRGECO, development of a silicone-based dural substitute to prevent adhesion of the healing dura to the adaptation of a 2P microscope to swine, and optimization of image acquisition protocols to reduce movement artifact and achieve images with sub-cellular resolution of neurons in vivo. Building upon work testing various strains of adeno-associated virus (AAV) expression in the swine brain^9^, we present a robust AAV transduction protocol, a synthetic cranial imaging window, and a 2P imaging system. These comprise a powerful toolset for testing hypotheses prior to embarking on human clinical trials to treat neurological diseases. As an example of the implementations of these technologies, we measure intracellular chloride ([Cli]), intracellular calcium ([Ca]), and extracellular chloride \ ([Cl_o_],) during epileptogenesis following traumatic brain injury in Yucatan minipigs.

## MATERIALS AND METHODS

A series of castrated, male, Yucatan swine (Sinclair Bio Resources LLC, Windham, ME; 5 months of age, 18-20 kg; n = 17) were used to optimize and refine techniques to measure [Cl_i_], [Cl_o_], and [Ca^2+^]. Four pigs were used to optimize AAV transduction and protein visualization via 2P imaging 1 month after AAV injection. After optimization, 11 pigs were used where [Cl_i_,], [Cl_o_], and/or neuronal [Ca^2+^] were successfully imaged with 2P imaging. Two swine (18 months of age, 76 - 84 kg) had received cortical impact 12 months prior to imaging. One pig had developed post-traumatic epilepsy; both were imaged to determine Cl_o_^10^. The cranial imaging windows were observed in 2 pigs 3.5 month after installation. All protocols were approved by the Institutional Animal Care and Use Committee of Massachusetts General Hospital and the Animal Care and Use Review Office of the Department of Defense. This manuscript describes the development of the technique. Here we describe the problem-solving methods and the techniques that were successful. We will report the effect of cortical impact on [Cl_i_] and [Cl_o_] in a separate manuscript.

### AAV injection surgery

Pigs were sedated with Telazol (2.2-4.4 mg/kg, intramuscularly, IM), xylazine (2 mg/kg, IM), and atropine (0.03 mg/kg, IM) and transported to the operating room. The pig was anesthetized with isoflurane and oxygen delivered via nose cone. An ear vein intravenous (IV) catheter was used to deliver cefazolin (25 mg/kg, IV), saline (2-4 mL/kg/hr), and propofol (as needed; 1 mg/kg). Swine were anesthetized with isoflurane and intubated and mechanically ventilated with oxygen. Tidal volume was adjusted to maintain a peak inspiratory pressure of 20-25 mmHg. End-tidal CO_2_ was maintained between 35 and 45 mmHg. Core body temperature was measured via a nasal or rectal probe and maintained at 37-39°C using a heating pad and Bair hugger forced air blanket. Blood pressure was measured via a cuff on a hindlimb and mean arterial pressure was maintained above 45 mmHg. Phenylephrine (25 – 60 µg/kg, IV) was administered for hypotension. End-tidal CO_2_, oxygen saturation, blood pressure, heart rate, and core body temperature were monitored and recorded every 5 minutes.

The pig’s head was secured to prevent movement and to mount the injector device. A human neurosurgery stereotaxic halo device (Cosman-Roberts-Wells) was tested first, but the pig’s head did not fit well, there were sharp pressure points, it lacked a maneuverable location to mount the microinjector pump, and it was not readily adjustable. Therefore, a stabilizing device was designed to fit the swine head and was constructed out of an aluminum optical breadboard and stabilizing rods (Thor Labs, Newton, NJ) and was secured to the operating room (OR) table (**Figure 1**). Two vertical rods were placed beside the neck on either side, with a horizontal rod connecting the two. Two pressure infusion bags were placed in between the two vertical rods and the pig’s head and inflated to further secure the head from movement without creating sharp pressure points. The pig’s legs were tied to the OR table. An OR table strap was placed over the pig’s upper back, and another behind the pig’s posterior to further secure the pig and prevent sliding (**Figure 1B**) while not impeding lung expansion.

**Figure 1.**
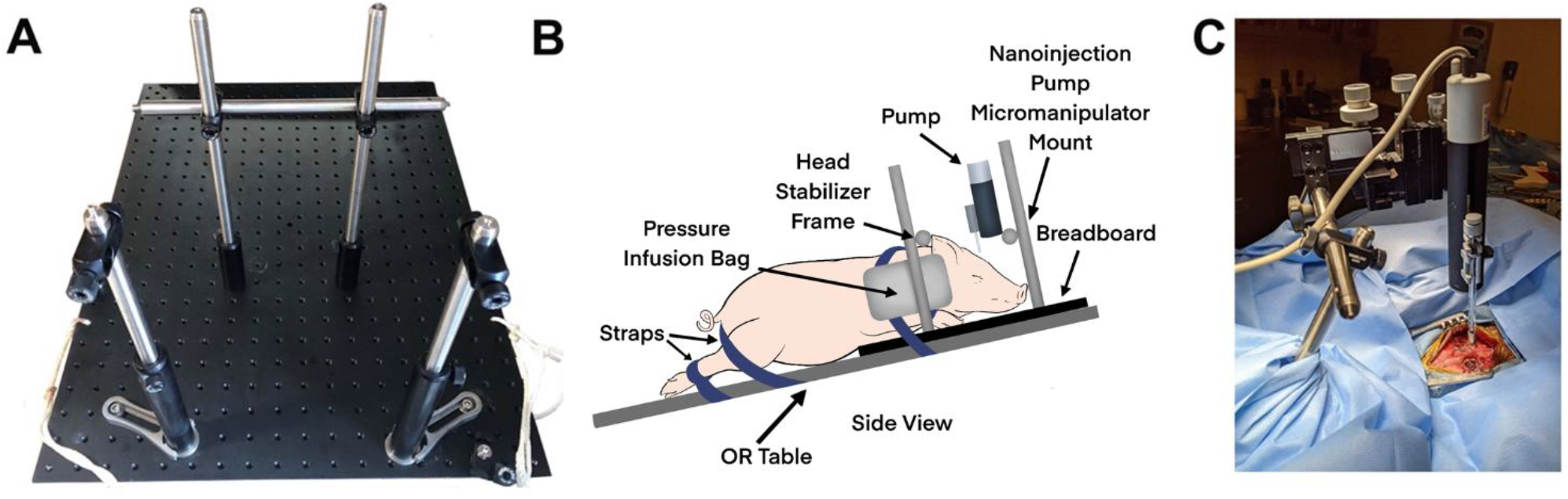
Surgery for cortical AAV injection and installation of the cranial imaging window. **A**. The pig’s head was secured in the breadboard with two vertical bars next to their snout and the horizontal bar pressing on the dorsal portion of its snout. Pressure infusion bags were inflated between the pig’s head and the back two vertical bars with the area between the chest and front legs pressed into the vertical bars. The pig’s front legs were tied to the board and the board tied to the OR table. Once the pig was draped, a sterile bar was placed horizontally between the back two vertical bars. **B**. Schematic depiction of the pig in position for AAV injection and imaging port implant. The pig was placed at a 30-45° angle to encourage the brain to fall away from the dura for implantation of PDMS avoiding cortical damage. **C**. The sterile cross bar added to the vertical bars after draping the animal. Then, the sterile injector and manipulator were mounted to the cross bar.

Fifteen minutes prior to fist incision, buprenorphine (0.02 mg/kg, IM) was administered. The head incision site was clipped and prepped using 70% alcohol, betadine, and chlorhexidine, and draped with antimicrobial incise drape (Ioban, 3M) and a large split-sheet drape and small drapes around the vertical bars anchored to the bread board. Bupivacaine (1.5-2.5 mg/kg, subcutaneously) was administered at the incision site before skin incision. The first incision was made running down the sagittal midline of the skull from above the snout to the crown of the head. The skin was detached from the periosteum to expose the skull. The sagittal and coronal sutures were located, and a Hudson drill was used to make a burr hole along the right coronal suture lateral to the sagittal suture over the rostral gyrus. A dural separator was used to detach the dura from the inside of the skull. Bone rongeurs were then used to expand the burr to approximately 2 cm in diameter. To minimize bleeding, electrocautery was used on the scalp and bone wax was applied to the edge of the skull. The dura was cut in a cruciate manner.

After the skull and dura were opened, the sterile cross bar was attached to the stabilizing device that was secured to the pig’s head (**Figure 1C**). The sterile manipulator and microinjector syringe pump (World Precision Instruments; Sarasota, FL) were attached to the cross bar of the stabilizing device (Figure 1A). The microinjector pump controller was placed on a non-sterile table next to the pig.

Vectors were synthesized by the Gene Transfer Vector Core facilities of the Schepens Eye Research Institute at Massachusetts Eye and Ear Infirmary, by the Penn Vector Core of the University of Pennsylvania, or by Addgene. Injections of AAV for SuperClomelon (SClm-AAV) with TurboRFP and/or CAG-tdTomato and/or jRGECO and/or nuclear localized green fluorescent protein mixed with 0.04% Trypan blue dye as a fiducial marker were injected into the cortex. AAV’s with the same wavelength of fluorescence were not injected into the same burr hole. AAV’s for TurboRFP, CAG-tdTomato, and nuclear localized green fluorescent protein were used as positive controls in the event that they were more robust than the target SClm-AAV or jRGECO. Virus was injected in 3-5 sites in each burr hole at a rate of 10,000 nL/min and approximate depth of 1 mm in locations not covered by cortical blood vessels. The location of each injection was mapped in relation to the array of cortical blood vessels. In one pig, injections at a depth of 2 mm or greater resulted in lesion and destruction of the cortical white matter. Slower injection rates such as 1,000 – 5,000 nL/min appeared to be so slow that the needle might have caused damage due to movement.

The anticipated shift in the skull over the following month was accommodated by injecting toward the midline of the pig’s head within the burr hole. An injection of SClm-AAV of 8.9 × 10^13^ gene copies/mL resulted in successful widespread cortical expression. Concentrations of 1 × 10^13^ gene copies/mL or below was not successful in transducing the cortex; however, however, the animal that received this dose also developed dural adhesions due to lack of a fully developed cranial window technique, so the lack of visualization of AAV-delivered gene products may not have been limited by low titer alone. In subsequent subjects, the order was switched such that the cranial imaging window was inserted first and then AAV was injected through the cranial imaging window for logistical reasons.

### Development of the cranial imaging window

To prevent dural adhesion to the cortex, the edges of the dural flaps were cauterized and initially, a Otologic Repair Graft (Circular 0.6 cm Laminated Extracellular Collagen Matrix, Cook® Biodesign®, Bloomington, IN) was placed under the edges of the dura to prevent the cortical surface from adhering to the healing dura. This was not successful, potentially due to rapid absorption. When the cortex is not protected from the healing dura, the dura adheres to the surface of the cortex and cannot be removed without removing cortical tissue and inducing trauma. We therefore synthesized and tested various types of polydimethylsiloxane (PDMS) membranes to act as a permanent barrier between the surface of the cortex and the healing dura.

Minimal clearance between the cortex and the dura made insertion of a PDMS membrane under the dura technically difficult. Strategies to allow the brain to shrink back from the dura allowed insertion of the PDMS without causing cortical damage. Two techniques were employed. The OR table was tilted 30-45 degrees such that the pig head was elevated above the body with the dorsum of the skull horizontal (**Figure 1**) adapted from Vuong et al.^11^ Additionally, ventilation was increased to reduce end tidal CO_2_ to 33-35 mmHg. A 1 mmHg reduction of PCO_2_ is associated with a 3-4% decrease in blood flow^12^. With these measures we found in unnecessary to employ additional chemical means (pentothal, mannitol, furosemide) to reduce the pressure of the brain on the dura as used in NHP^13^.

The dural substitute was rinsed in sterile saline, rolled, and gently placed under the dura and slowly encouraged to unfold. After unfolding, the dural substitute was centered such that the dural substitute overlayed the entire exposed cortex and the edges of the dural substitute extended under the dura for at least 5-10 mm (**Figure 3A**). The dural flaps over the PDMS were opened further as wide as possible and additional cautery was performed as needed and the underside of the dura was glued to the PDMS using Histoacryl skin adhesive (McKesson, Irving, TX) creating a seal. PDMS was inserted under the dura of both burr holes.

With the first iteration of the dural substitute, we attempted to use a PDMS membrane with an attached silicone ring that would line the burr hole and secure the PDMS, because the dura would grow around the silicone ring (**Figure 2B**) similar to that described by Arieli et al.^13^. However, we found this version of the dural substitute difficult to place underneath the dura without inducing trauma to the cortex and/or cortical blood vessels. We then removed the silicone ring and used a flat membrane only as a dural substitute (**Figure 2C**). The flat membrane-only version was easier to place underneath the dura, but we found it would easily relocate if not secured. The dura was secured to the PDMS with suture but the suture ripped through the PDMS or increased the risk of bleeding as the suturing needle can tear cortical veins. In our next attempt, two or four silicone straps were embedded into the membrane (**Figure 2A**). These straps were then glued to the skull. This version was easy to insert under the dura and worked well to secure the PDMS but would rip/detach during cortical impact over the PDMS. The final version we used was the plain membrane disk which we secured by gluing to the dura (**Figure 2D**). Ridges on the dura helped identify the center of the disk for placement. This version was relatively easy to place, had a low rate of complications, allowed us to use our mechanical cortical impact device over inserted PDMS, and kept the dural substitute intact after cortical impact. Further refinements of the cranial window and easier access at 2P imaging included insertion of a polypropylene ring not attached to the PDMS. The rigid ring served to reduce skull and tissue remodeling above the PDMS site allowing easy access and a pristine compartment closed off by the body at 2P imaging approximately 1-3.5 months later (**Figure 3**).

**Figure 2.**
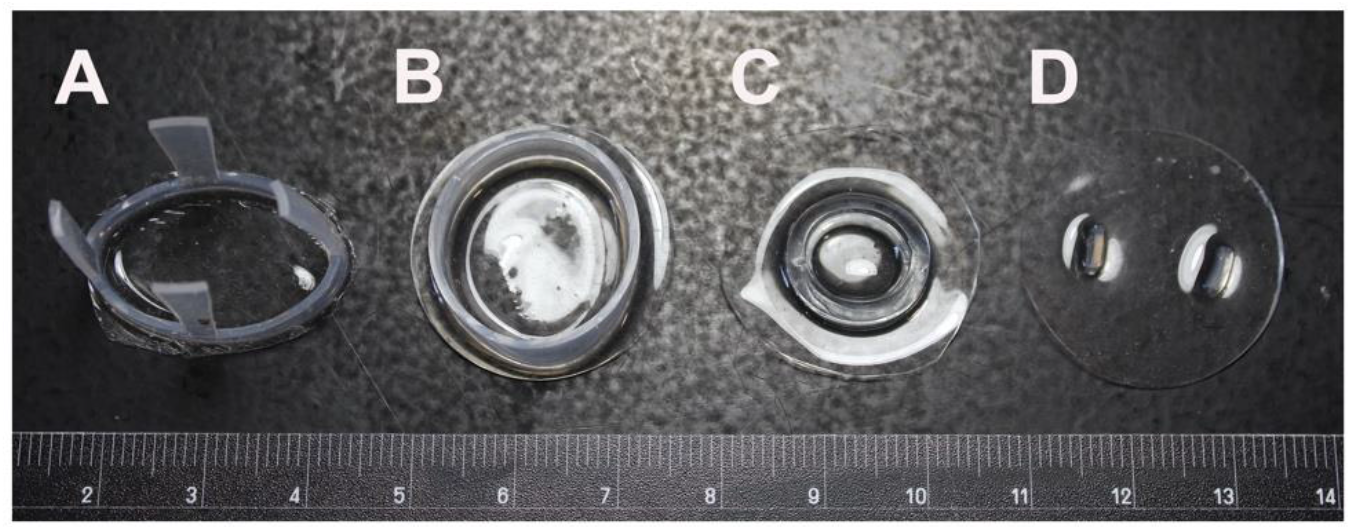
The evolution of the dural substitute. A. PDMS with silicone straps worked well but prevented later cortical impact. **B**. PDMS with “top hat” portion to fill the edges of the burr hole. This version was difficult to place without causing cortical trauma. **C**. PDMS with flat silicone ring. The ring didn’t provide much advantage and limited the area for 2P imaging. **D**. Flat PDMS with ridges to identify the center of the PDMS. This iteration was relatively easy to install, allowed centering of the PDMS, and could withstand injections and cortical impact.

**Figure 3.**
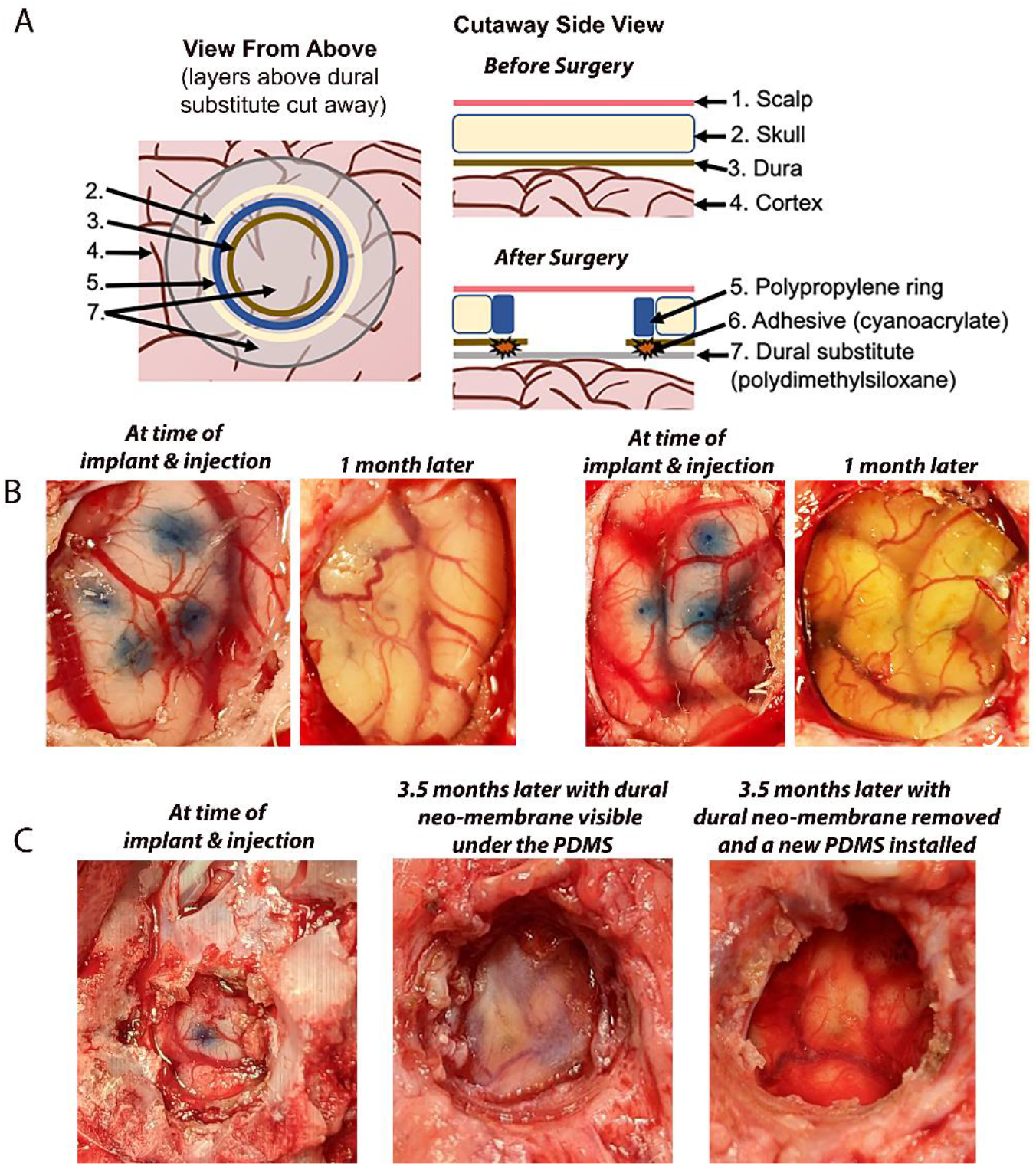
Cranial imaging window. **A**. The cranial imaging window system from the top and the side that prevented the adhesion of the healing of the dura onto the cortex that obfuscated the surface of the cortex as well as tissue growth from the burr hole of the skull. **B**. Two burr holes displaying the PDMS overlaying the cortex immediately after AAV injection and the pristine window 1 month later. **C**. The cranial window at installment, after 3.5 months with dural neomembrane obscuring the cortex, and immediately after dural neo-membrane removal and PDMS re-installation. This iteration of the cranial window had an adjacent subdural electrode strip that entered posterior to the imaging window.

Upon completing the injections and dural implant insertion, the cortex was thoroughly flushed with sterile saline and the incision was closed with interrupted subcutaneous sutures using 2-0 PDS suture for the subcutaneous layer followed by a running subcuticular stitch using 3-0 monocryl. The incision was cleaned and skin adhesive was applied. Once the pig was stable, anesthesia was lightened and the pig was encouraged to breath on its own and was extubated. Buprenorphine was administered (0.025 mg/kg, IM) and fentanyl transdermal patches (1-4 µg/kg/hr for 72 hours post-operatively) were placed on the lower back for pain management. The animal was then transferred to the animal facility and monitored until ambulatory. The animals were observed twice a day for three days post-operatively and five times a week until fully healed from surgery.

### Large animal 2P microscope

The 2P microscope for imaging swine was designed with four key specifications: 1) ability to accommodate large animals with considerable flexibility, 2) intensity and fluorescence lifetime imaging without switching detection hardware and scan path, 3) externally triggered high speed volumetric scanning, and 4) low-cost construction from off-the-shelf parts. The overall design of the microscope was based heavily on the previously published open source TIMAHC (Two-photon Imaging that is Modular, Adaptable, High-performance and Cost-effective) design^14^ with exceptions detailed below.

#### Accommodation of large animals

To provide a large open space for staging of the pig’s head, the microscope was built using a gantry design, where the TIMAHC’s vertical beam containing the scan head and detection hardware was mounted to a girder (**Figure 4**). The girder was affixed between two vertical beams, which were mounted on a pair of translating stages (XYR8080, Dover Motion, Dover, MA): one motorized and one passive. The objective lens and detection module were mounted to a motorized stage (Sutter-1z) for motorized focus adjustment. Together, the gantry design provided a large berth for the animal’s head, the motorized stages provide approximately 150 × 150 mm of microscope travel in the × and y directions and 25 mm of travel in the z-direction, with additional manual height adjustment of up to 50 cm in the z-dimension.

**Figure 4.**
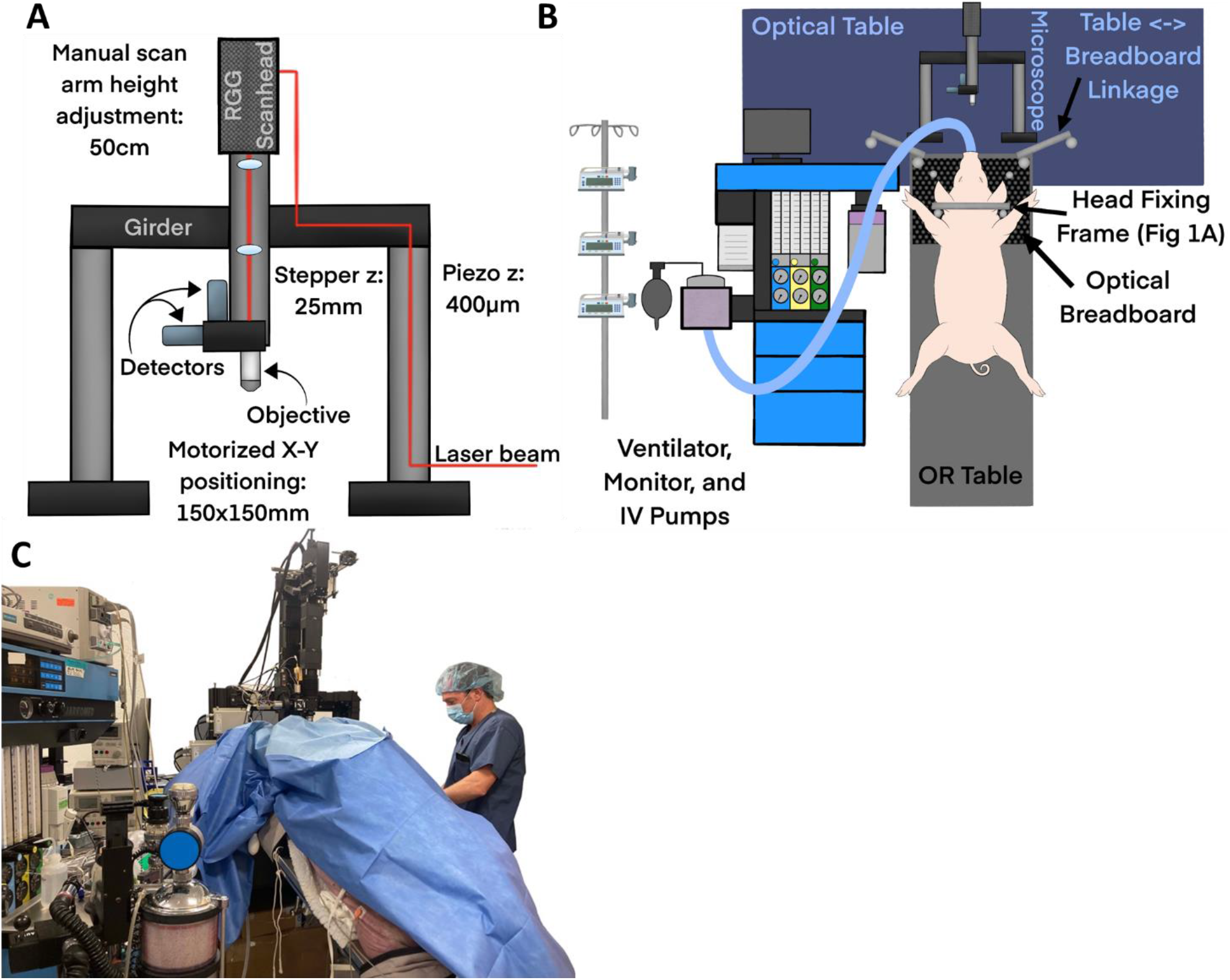
2-photon rig and anesthetic support set up. **A**. Schematic depiction of key elements of the microscope body, highlighting the large range of motion for accommodating a large animal. **B**. Schematic of room layout during an imaging session. **C**. Photograph of an imaging experiment with an 84 kg pig under the 2-photon microscope. The surgery table was resting on and mechanically coupled to the optical table. The ventilator is placed perpendicular to the optical table to the left of the pig with the ECG, gas analyzers, physiology monitor (not shown) and IV pumps on the ventilator (scale: human = 1.7 meters).

#### All-digital fluorescence lifetime and intensity measurement

Conventional 2P imaging involves scanning a focused laser across a sample, while integrating the output of an emitted light detector (typically a photomultiplier tube) for a pre-determined pixel dwell time to measure the fluorescence intensity of each pixel. Fluorescence lifetime imaging (FLIM) characterizes the timing of fluorescence emission following excitation. Because of special requirements for quantifying the timing of emitted photons with nanosecond precision, FLIM detection is commonly performed with an independent set of detectors and detection electronics. In the system presented here, light was detected with hybrid photodetectors (R11322U-40, Hamamatsu USA, Bridgewater, NJ), which have a low transit time spread, producing an output pulse of approximately 1 ns for each photon detected. A high-speed digitizer (High-speed vDAQ, MBF Bioscience, Williston, VT) was used to digitize this signal at 2.56 GHZ with a sampling clock that was frequency multiplied from (and hence phase-locked to) the laser pulse. Together with firmware from Scanimage (MBF Bioscience), this hardware combination enabled simultaneous photon-counting based intensity imaging and digital FLIM in a single light path, simplifying image acquisition protocols. The vast majority of fluorescent proteins and commonly used organic dyes have fluorescence emission lifetimes between 1 and 8 ns. For fluorophores with a lifetime in the range of 1-8 ns, digital FLIM provides accurate and substantially more photon-efficient lifetime measurements due to a lack of detection “dead time” inherent to time-amplitude converter based FLIM system (e.g. Fig. S1 shows a digital FLIM-measured lifetime histogram for sodium fluorescein, an organic dye with a published of ∼4.1ns). Thus, lifetime images can be acquired in a shorter period of time, which is advantageous for large animal imaging.

**Supplemental Figure 1.**
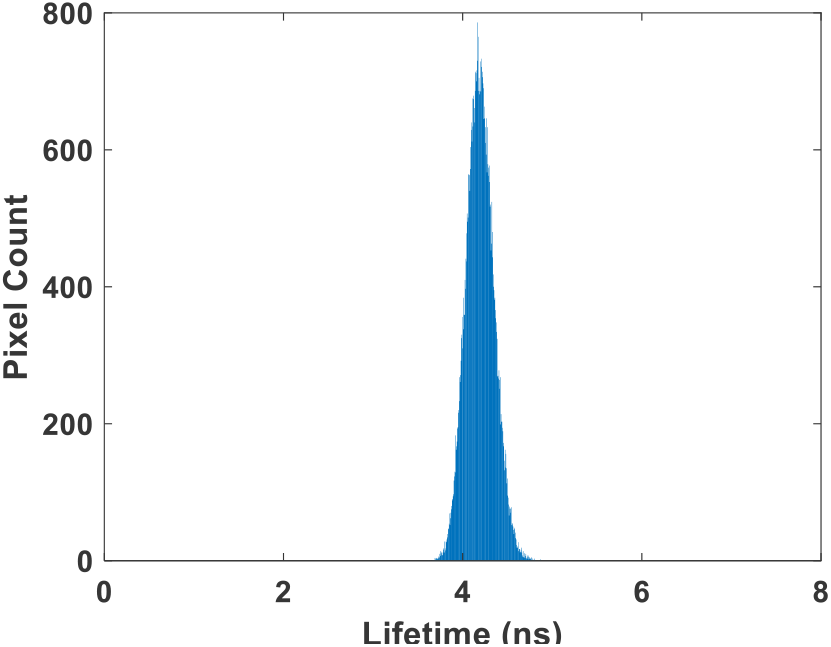
Fluorescence lifetime histogram of sodium fluorescein. Our digital FLIM system reported a mean lifetime of 4.197 ns, with a standard deviation of 155ps.

#### Triggerable high-speed volumetric scanning

When imaging a > 80 kg animal, the motion artifact is significant. The sources of motion artifact (lungs, heart) scale proportionally with animal size, while the objects being imaged (neurons, glia, extracellular matrix) remain small. As described below, the respiratory and cardiac movement artifacts were mitigated with brief, induced apnea and electrocardiogram (ECG)-triggered image stack acquisition. Triggering image acquisition on the QRS complex means that all images will be acquired in the same phase of the cardiac cycle, which enables post hoc motion correction algorithms and averaging of multiple acquisitions. However, it requires that image stacks be acquired in the window of time *between* systoles. The volume acquired was maximized using high speed resonant scanners (RMR scanner, MBF Biosciences) to scan in the x-y plane and a piezoelectric focusing device (P-725 with E-665, Physik Instrumente) to move the objective in the z plane. Together, this system can scan ten 512 × 512 pixel slices (a 600 × 600 × 400 µm volume) in < 400 ms.

#### Low-cost construction from off-the-shelf components

With the few notable exceptions listed above, the microscope was constructed of readily available parts detailed in Rosenegger et al.^14^ During construction in 2020, all parts, including the laser and air table were obtained for a total cost of < $250K.

### In vivo imaging of the transduced cortex

At 3-5 weeks after AAV injection, pigs were imaged using a 2P microscope custom built to accommodate the swine. At 12 months after cortical impact or sham surgery, an additional cohort of pigs was acutely injected with a dextran-linked fluorophore (using the injection procedure described above) and imaged^15^.

Moving the large pigs (76-84 kg) to the campus where the microscope was located required a specialty moving company. A rigid stretcher on wheels was used to move the pig to the OR table requiring at least 4 staff to manually lift and position the pig.

To access the cranial imaging window, pigs were sedated, instrumented, and positioned in the stabilization device as described above. General anesthesia was maintained with isoflurane titrated to 1-2% mixed with medical air. An ear vein IV was used to deliver a dose of cefazolin (25 mg/kg, IV) and a continuous infusion of saline (2-4 mL/kg/hr IV); pigs with hypotension received a continuous infusion of dexmedetomidine (1-2 µg/kg/hr, IV). The head incision area was prepped for surgery as described above except that only one layer of drapes was used as it was a non-survival surgery. Prior to incision, the OR table was tilted to decrease intracranial pressure. A T-incision was made on the scalp and the skin was detached from the skull. Mechanical ventilation was adjusted to reduce end tidal CO_2_ to 33-35 mmHg. The underlying scar tissue was detached from the skull and the burr holes were carefully resected being careful not to puncture the dural substitute. Resection was made easier with the dural retention device in place, which resulted in encapsulation of the PDMS and clear fluid filling the burr hole above a pristine cortex. The periosteum was cleared and a well to hold saline was created by constructing a wall of Maxcem Elite dental acrylic (Kerr Corp., Brea, CA). The well encompassed both burr holes and was filled with saline allowing a meniscus to form between the microscope objective and saline. In the later subjects, a meniscus was achieved without the dental cement well.

The OR was arranged so that the ventilator was adjacent to the optical anti-vibration table (**Figure 4B**) and the initial surgery was performed with the pig’s head away from the optical table. Once the cranial imaging windows were opened and the well was constructed, the table was rotated so the head was towards the microscope and then the OR table rolled forward to make contact with the optical table. On the initial attempts, the head was placed on a mount on the optical table, but this was difficult to secure and instead, the pigs head was secured in the breadboard stabilizing device onto the OR table and the OR table was rested on the optical table for imaging sessions thereafter. The objective of the microscope was sterilized with topical 70% alcohol for at least 30 seconds.

### Reducing Movement Distortion

The pig secured to the breadboard was secured to the optical imaging table to prevent artifact movement. To remove breathing artifact during image acquisition, the pig was switched to from room air to 100% oxygen, paralyzed with boli of vecuronium (1 mg/kg) every 20-30 minutes, and the ventilator turned off. Each episode of breathing cessation lasted 2-2.5 minutes and peripheral oxygen saturation was minimally affected (above 93%).

Movement distortion from the cardiac cycle was reduced by timing laser pulses to trigger between systole via the ECG^16^. A 5-lead ECG was connected to a monitor (Datex-Ohmeda Cardiocap™ 5; Datex-Ohmeda, Madison, WI) and the “Defib Sync” QRS detection transistor-transistor logic output was connected to the microscope control hardware for triggering image acquisition phase-locked to the ECG.

### Cortical Impact

Upon completing the pre-injury chloride imaging, pigs received a cortical impact to one hemisphere through the cranial imaging window, the other cranial imaging window serving as a control. The cortical impactor stand was secured over the burr hole with its three feet securely planted between the skull and over the dural substitute flaps. The three support screws were firmly secured to the skull^17^. The cortical impact device was then screwed into the stand until the impactor tip (1.07 cm in diameter) was just touching the surface of the dural substitute. The indenter was deployed (over 4 ms) over the dural substitute with an indentation velocity of 1.5 - 1.7 meters/second mimicking a closed dura injury, as we have previously described^17^. The indenter and stand were then carefully removed. Any resulting blood was gently flushed with saline until the cortex was visible. If blood pooled under the dural substitute, saline was flushed using a 25-gauge needle through the dural substitute. After cortical impact, the pig was then imaged again through the intact PDMS.

### Revision of the cranial imaging window

As the PDMS was flexible and the retention ring assembly was subcutaneous, adjusting the PDMS and burr hole if needed was possible. In one instance, cortical impact through the PDMS tore the PDMS and in another instance, flushing the surface of the cortex with a saline with a needle through the PDMS ripped the PDMS and in these cases, a new membrane was inserted and secured. If the injection sites on the cortex were under the skull due to skull remodeling, the edges of the burr hole could be extended and the PDMS re-secured.

### MRI

After acquiring the pre-injury and 1hr post injury 2P images, a subset of pigs received MRI. Those results will be reported in a separate paper. The dural substitute was made of MRI compatible material and was not removed prior to MRI and did not cause artifact.

### Transcardiac Perfusion

After imaging was complete, the pig was euthanized via exsanguination by transcardial perfusion with 0.9% saline and 10% formalin and the brain was collected. The brain was post-fixed in 10% formalin for 5 days and then moved to PBS. The brain was cut into 5 mm slabs, paraffin embedded, 10 µM sections were made, and then de-waxed. The expression of SClm was observed and photographed on a fluorescent microscope. The pattern of expression of SClm in the gyri were mapped by hand and diagrammed.

## RESULTS

Rapid and shallow (1mm depth) injection of SClm-AAV and other AAV vectors successfully transduced of the cortex with wide-spread SClm expression without damage to the cortex (**Figure 5)**. Histology revealed that the expression of SClm was prominent through Layers 2/3 of the cortex and even down to layer 6 and into the gyral white matter (**Figure 5**) deep in the base of the gyri. With the cranial imaging window keeping the surface of the cortex pristine and the 2P microscope designed to accommodate significant respiratory and cardiac movements of swine up to 84 kg, subcellular structures could be resolved, and image quality was sufficient to enable ratiometric assessment of [Cl_i_] (**Figure 6**) and fluorescence lifetime-based [Cl_o_] (**Figure 7**).

**Figure 5.**
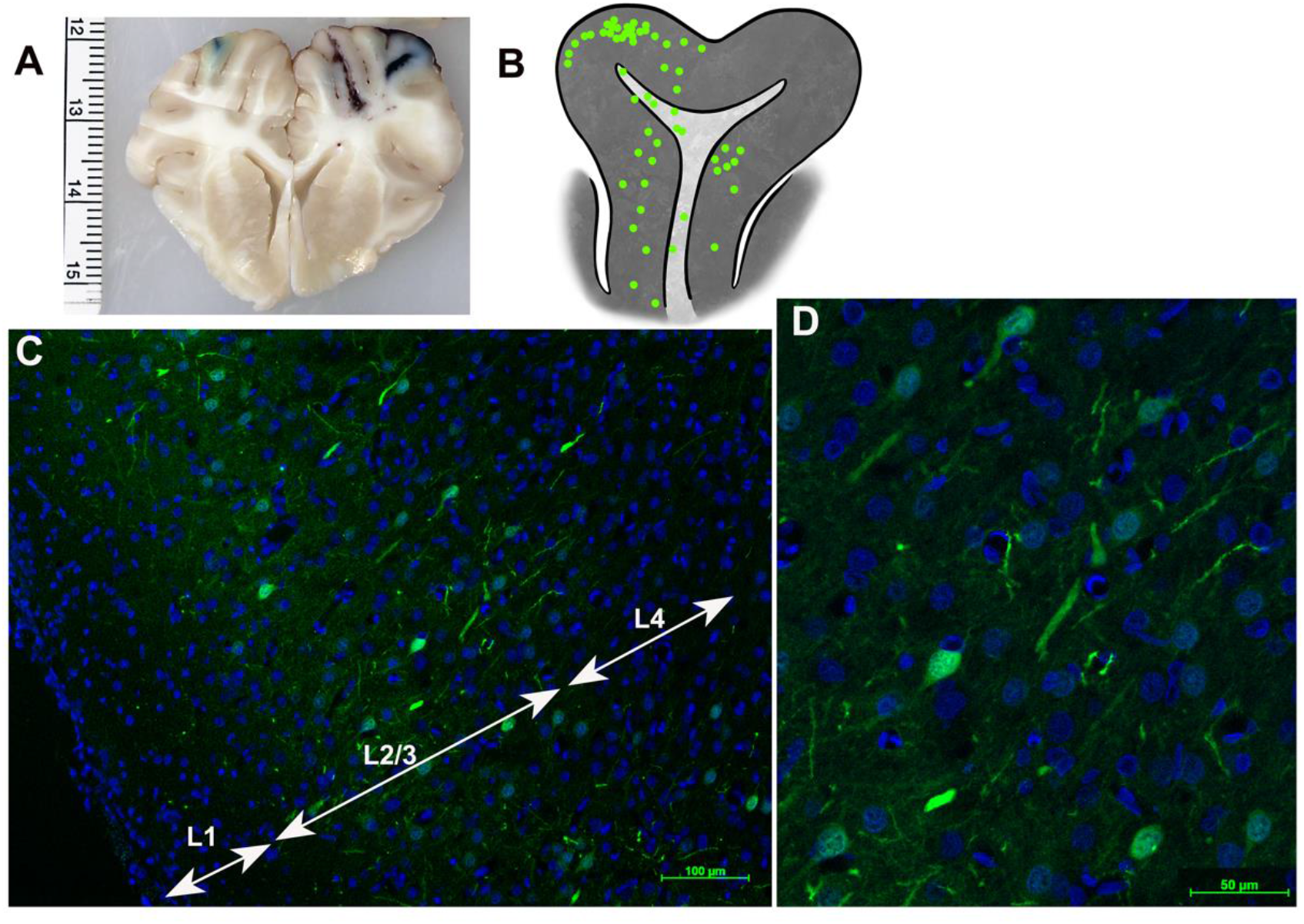
Transduction of the cortex with SuperClomeleon. **A**. Coronal slab of brain with bilateral injections of SuperClomeleon AAV into the rostral gyrus, 5 weeks after injection and 5 hours after unilateral cortical impact (right hemisphere). As expected, the cortical impact site is moderately swollen with a thin subarachnoid and with intraparenchymal hemorrhages and the injection site visible (blue dye). On the left cortex, the injection site (blue dye) remains pristine following AAV injection and after 2 hours of 2P imaging. **B**. Schematic demonstrating the distribution of SuperClomeleon^+^ neurons in the rostral gyrus down to the sulci. SuperClomeleon^+^ neurons were dense in Layers 2/3 of the gray matter, were dispersed through all layers to Layer 6, and were in the gyral white matter, perhaps expressed by interstitial neurons. **C**. Low power (10x) photomicrograph of SuperClomeleon^+^ neurons (green) with nuclei (DAPI, blue; scale bar = 100 μM). SuperClomeleon^+^ neurons are dense in Layer 2/3^18^ and dispersed in Layer 4. **D**. Higher power photomicrograph (20x; scale bar = 50 μM) of Layer 2/3 in Panel **C** demonstrates SuperClomeleon expression in pyramidal neuron cell bodies and neurites.

**Figure 6.**
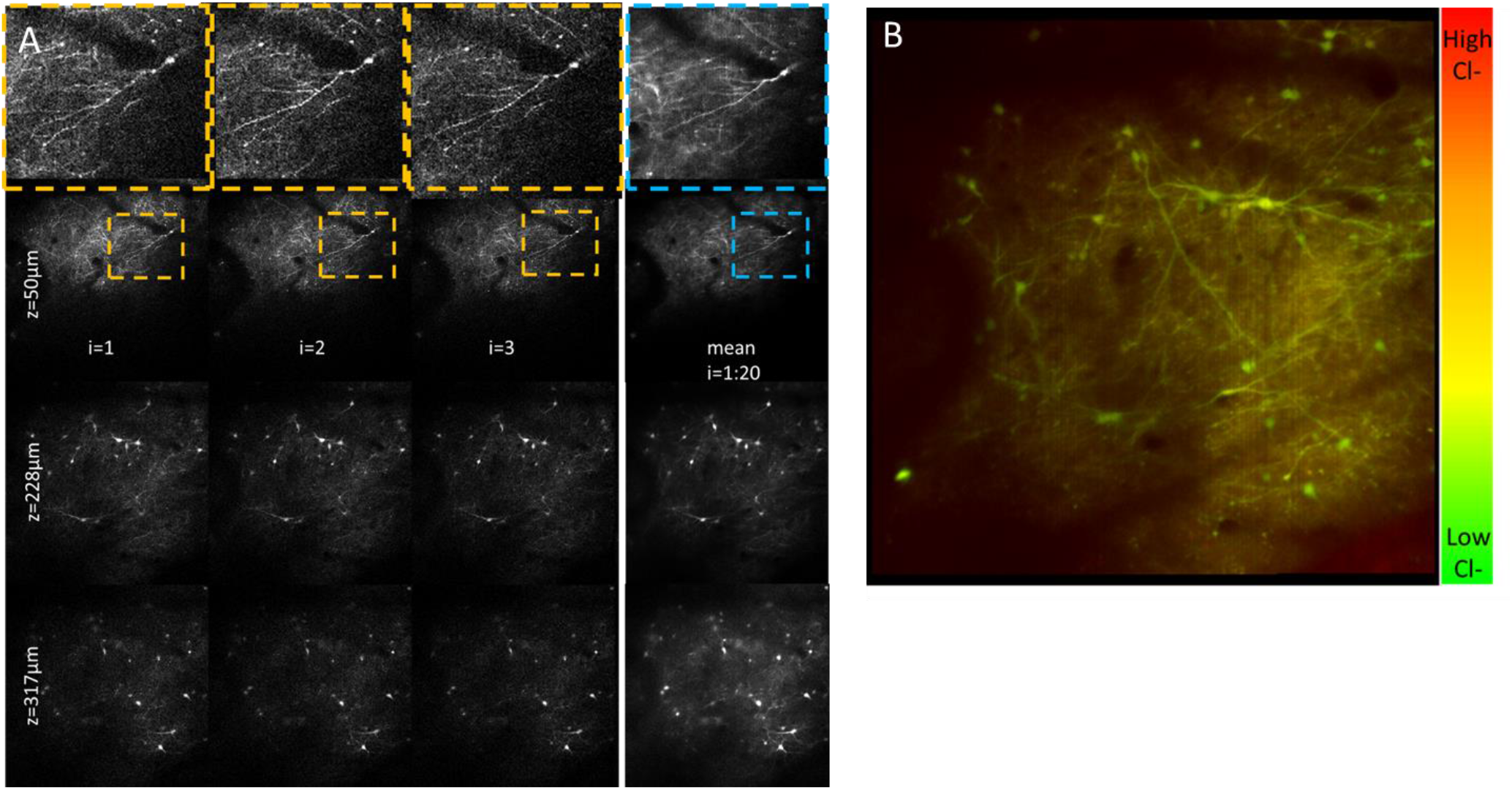
Imaging in vivo intraneuronal chloride of the cortex. **A**. The heartbeat triggered acquisition of a fast z-stack spanning 600 × 600 × 400 µm provided stable images of the fluorescent protein-transduced neurons *in vivo*. Because these volumes were acquired between heartbeats, individual iterations (first 3 columns) could be averaged together to increase dynamic range and decrease shot noise in the final image stacks (4^th^ column). **B**. Merging the cyan and yellow fluorescent protein signals from SuperClomeleon provides a pseudo-colored imaged of intracellular chloride.

**Figure 7.**
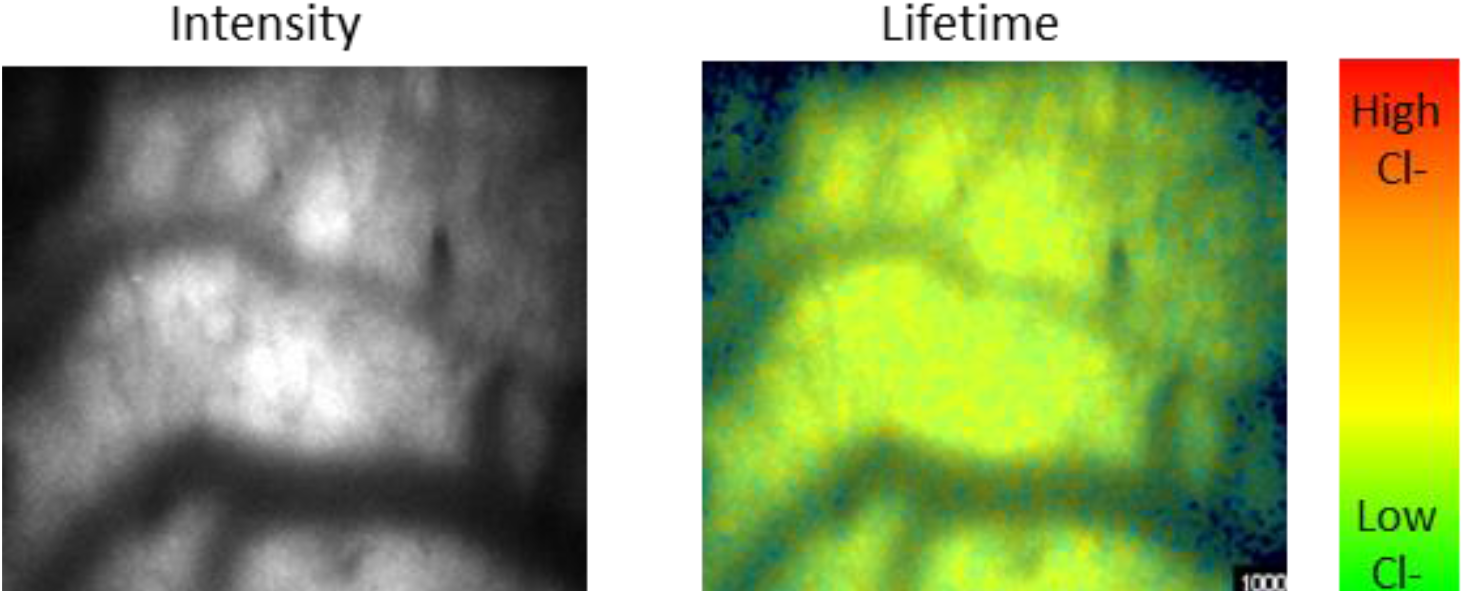
Imaging in vivo extracellular chloride of the cortex. After injection of a 10 kD dextran-linked chloride sensitive organic dye, fluorescence lifetime images were acquired using the heartbeat-triggered volume acquisition described above. This produced high resolution intensity (left) images and lifetime (right) images that can be used to quantify extracellular chloride.

Serial imaging appears to be possible because the cranial imaging window was functional and revisable for up to 3.5 months and the cortex of the imaging site pristine after several hours of imaging (**Figure 5**). At 3.5 months an opaque dura neo-membrane was observed in one cranial imaging window below the PDMS but not adhered to the cortex. The dural neomembrane was easily removed and a new PDMS was installed. The dural substitute was also thin enough that the micro syringe was able to inject through it while being robust enough that it did not tear during cortical impact at the time of chloride imaging.

### Morbidity and Mortality

No swine had an infection from the surgery or from the fully subcutaneous cranial imaging windows. ^11^ One pig developed neurologic symptoms and was euthanized early after the placement of cranial imaging window. The PDMS iteration in this case had a silicone ring/ “hat” attached to line the burr hole and more difficult to place. There was a large space-filling hemorrhage under the PDMS membrane in an animal that had previously placed skull screws for EEG^19^ perhaps limiting movement and accommodation of the brain to the hematoma. Placing the PDMS membrane does present a risk of laceration of cortical veins and re-bleeding after skin closure. We reduced the risk of hemorrhage by placing a PDMS without the attached ring and ensured that all bleeding had ceased before closing the incision.

## DISCUSSION

Swine provide advantages to NHP insofar as cost and public perception but have major anatomical and behavioral differences that required creation of new approaches. Here we present adaptation of AAV cortical transduction, a cortical imaging window, and in vivo, 2P imaging to swine ranging from 20-84 kg. Our goal was to make a cranial imaging window that allowed repeat access to the transduced cortex without dural adhesion or formation of a dural neo-membrane with a low risk of infection. Additionally, for our application of studying traumatic brain injury, the cranial imaging window would accommodate cortical impact and could be installed without administration of pharmaceutical agents to withdraw the brain from the dura^13^ that may also affect the response of the brain to cortical impact.

We chose recombinant AAV-9 because previous work in swine demonstrated that recombinant AAV-9 with GFP produced the best expression compared to other serotypes^9^. We chose intraparenchymal injections vs. administration via the cisterna magna to more efficiently transduce the cortex of interest, improving expression levels with a lower volume of AAV (thus, reducing cost). Qualitatively, it appears that the expression described here was localized to a radius of approximately 5 mm around the injection site, but otherwise similar to previously reported cisterna magna expression in density and depth^9^. Our findings may indicate that the virus easily spreads in the CNS regardless of route. Here, inherent limitations of 2P imaging, not recombinant AAV transduction, was the limiting factor for the depth of visualization. Visualization beyond layers IV/V of the cortex would require adding complexity to the imaging system such as a gradient index of refraction lens-based endoscope, microprism, or three-photon excitation.

There were two major challenges in translating viral transduction and fluorescence imaging techniques to swine: 1) swine do not accommodate head posts well and 2) swine have extensive accretion and remodeling of the skull. Unlike NHP and rodent species, swine are known to move their head against objects, have large neck muscles, a thick skull, and can damage any exteriorized equipment leading to destruction and/or infection. However, unlike NHP, swine do not pick at incisions. Strains of NHP are small, are housed for long durations, and can be trained to sit quietly for weekly servicing of their exteriorized ports with antibiotic washes and bacterial surveillance^11^. Conversely, restraint of swine is more difficult requiring anesthesia in a research setting and they are not typically kept for the long duration necessary for the training to achieve weekly image port servicing without restraint or anesthesia. We adapted a transparent PDMS membrane used in NHP to seal the dura and prevent formation of dural adhesion, but instead of exteriorizing it, we kept the entire assembly subcutaneous^13^. This strategy prevents the need for weekly servicing and prevents destruction of the window by swine-typical behavior. The subcutaneous strategy also likely reduced the possibility of infection. However, an exteriorized capped window could be trialed and the labor required/cost for weekly washing and culture compared to surgical incisions in a survival OR.

We observed that dural neo-membranes formed sometime between 1 month and 3.5 months in a subset of cranial imaging windows as has been demonstrated in NHP using voltage-sensitive dyes^13^. Removal of the neo-membrane without injuring the cortex was possible as demonstrated in NHP^13^. Our revisable cranial window was easily re-built after removal of the dural neo-membrane. Filling the window with antibiotics and agarose or fibrin was not necessary with our subcutaneous window. Indeed, even a subset of “permanent” windows in NHP also required re-building at these later imaging sessions^13^.

The vast amount of skull accretion and remodeling in swine presented another reason to make the cranial window easy to “edit”. The adult rhesus monkey weighs 8-11 kg. The vault of the skull is rounded and is 1.5 – 2.5 mm thick (skull: bodyweight ratio of 0.18). Though the Yucatan pig is domesticated, their skull has some characteristics similar to wild boar. The vault of the immature Yucatan is relatively flat compared to the Yorkshire strain and is even concave after maturity is reached. In the Landrace strain, the skull grows from 15 mm thick at 28 kg (skull: bodyweight ratio 0.5) to 22 mm thick at 70 kg (skull: bodyweight ratio: 0.3) while the size of the brain remains unchanged^8^. In the Yucatan, the nuchal crest located on the crown of the skull, which functions to anchor their powerful jaw muscles, can grow up to 30-50 mm thick. During development, sinuses grow over the top of the forehead of the skull and the angle of the brain changes in relation to the skull with the rostral portion moving ventrally. The thickness of the skull relative to bodyweight is 2-3 times greater in swine than NHP. Because of the skull remodeling, a revisable PDMS window allows adjustment of the window during skull accretion. Here, we found that if AAV was injected medially in the burr hole, the portion of the transduced skull remained visible as opposed to lateral injection sites that would end up under the remodeled skull 1-3.5 months later. Regardless, expanding the burr hole at the time of the imaging session to expose potentially buried AAV injection sites is feasible with the cranial window construct described here.

## CONCLUSION

Establishing swine as a gyrencephalic model for preclinical studies is an ongoing effort aimed at improving translatability of therapies for neurological disorders. Here we aim to facilitate the expansion of such studies by providing detailed protocols for two tools (AAV transduction and 2P imaging for large animals) that are widely used in preclinical studies and have a broad range of applications.

## Abbreviations

(NHP): non-human primates
(2P): 2-photon
(OR): operating room
(AAV): adeno-associated virus
(PDMS): polydimethylsiloxane
[Cl_i_]: intracellular chloride
[Cl_o_]: extracellular chloride
(SClm): SuperClomeleon
(TIMAHC): Two-photon Imaging that is Modular, Adaptable, High-performance and Cost-effective
(FLIM): fluorescence lifetime imaging

## ACKNOWLEDGEMENTS

This work was supported CURE Epilepsy based on a grant CURE Epilepsy received from the United States Army Medical Research and Materiel Command, Department of Defense (DoD), through the Psychological Health and Traumatic Brain Injury Research Program under Award No. W81XWH-15-2-0069 to KJS. Opinions, interpretations, conclusions and recommendations are those of the author and are not necessarily endorsed by the Department of Defense. In conducting research using animals, the investigator(s) adheres to the laws of the United States and regulations of the Department of Agriculture. It was also supported by NIH K01 HD083759, R01 HD09939, and a Claflin Distinguished Scholar award from Massachusetts General Hospital to BCB, and NIH R01 NS112538 to KPL.

The authors would like to thank Bryan Golemb, Andrew Ding, John Dempsy, Caroline Kaplan, Kevork Marco Rodriguez Hovnanian, Akhila Penumarthy, Alex Hone, who helped with the surgeries, anesthesia, perfusion, and brain collection. We thank the team at Knight Surgery for providing anesthesia support for the AAV injection surgery. We thank the CCM team and Titi Lamidi for the long-term care of the animals and organizing the shipping of large pigs to the campus where the microscope was located. We thank Dr. Andrea Slate and her team for her advice and assistance with post-surgical care. We thank Dr. Ann-Christine Duhaime for the recommendation of the dural graft that we tested. We thank Dr. Brittany Coats and Robert Metcalf at the University of Utah for assessment of biomechanical properties of the indentors.

